# Evolving a generalist biosensor for bicyclic monoterpenes

**DOI:** 10.1101/2021.08.20.457167

**Authors:** Simon d’Oelsnitz, Vylan Nguyen, Hal S. Alper, Andrew D. Ellington

**Affiliations:** Department of Molecular Biosciences, University of Texas at Austin, Austin, TX, 78712, USA; Freshman Research Initiative, University of Texas at Austin, Austin, TX, 78712, USA; McKetta Department of Chemical Engineering, University of Texas at Austin, Austin, TX, 78712, USA

**Keywords:** Biosensors, Protein Engineering, Directed Evolution, Terpenes, Metabolic Engineering

## Abstract

Prokaryotic transcription factors can be repurposed as analytical and synthetic tools for precise chemical measurement and regulation. Monoterpenes encompass a broad chemical family that are commercially valuable as flavors, cosmetics, and fragrances, but have proven difficult to measure, especially in cells. Herein, we develop genetically-encoded, generalist monoterpene biosensors by using directed evolution to expand the effector specificity of the camphor-responsive TetR-family regulator CamR from *Pseudomonas putida*. Using a novel negative selection coupled with a high-throughput positive screen (Seamless Enrichment of Ligand-Inducible Sensors, SELIS), we evolve CamR biosensors that can recognize four distinct monoterpenes: borneol, fenchol, eucalyptol, and camphene. Different evolutionary trajectories surprisingly yielded common mutations, emphasizing the utility of CamR as a platform for creating generalist biosensors. Systematic promoter optimization driving the reporter increased the system’s signal-to-noise ratio to 150-fold. These sensors can serve as a starting point for the high-throughput screening and dynamic regulation of bicyclic monoterpene production strains.

## INTRODUCTION

The rapid and facile chemical measurement of biologically produced compounds is crucial for the $14.4B high-throughput screening industry, especially when discrimination is needed for enantiomers or constitutional isomers^1^. In particular, there remains a pressing need for the development of high-throughput analytical methods for terpenes used commercially in the flavor, fragrance, cosmetic, and pharmaceutical industries. Although monoterpenes are traditionally sourced from plants, microbes are being increasingly engineered to provide a more reliable, pure and space-efficient means of production^2^, typically via heterologous expression of a pathway for the production of the common intermediate geranyl pyrophosphate (GPP) followed by a specific terpene synthase. Since most monoterpenes lack a chromophore and terpene synthases often nonspecifically produce a range of terpenes with the same exact mass, metabolic screening is typically limited to low-throughput gas chromatography-mass spectrometry.

Prokaryotic transcriptional factors can be repurposed as genetic biosensors for high-throughput chemical measurement and metabolic regulation, thereby leveraging specific protein-ligand interactions to selectively transduce chemical abundance into an easily quantifiable fluorescent output^3^ (**Figure 1a**). Genetic biosensors for monoterpenes would circumvent the aforementioned analysis challenges and enable high-throughput screening for improved strain variants. However, genetic biosensors are not widely used, largely because the range of natural compounds that can be sensed is far smaller than the repertoire of chemicals and metabolites that see industrial applications. Bioinformatic mining conjoined with high-throughput experimentation has accelerated the pace of natural biosensor discovery^4^, but only a few natural transcriptional regulators responsive to monoterpenes^5,6,7^ or derivatives thereof^8^ have been identified, and to date no genetic biosensors have been engineered or thoroughly characterized for monoterpene analysis.

**Figure 1.**
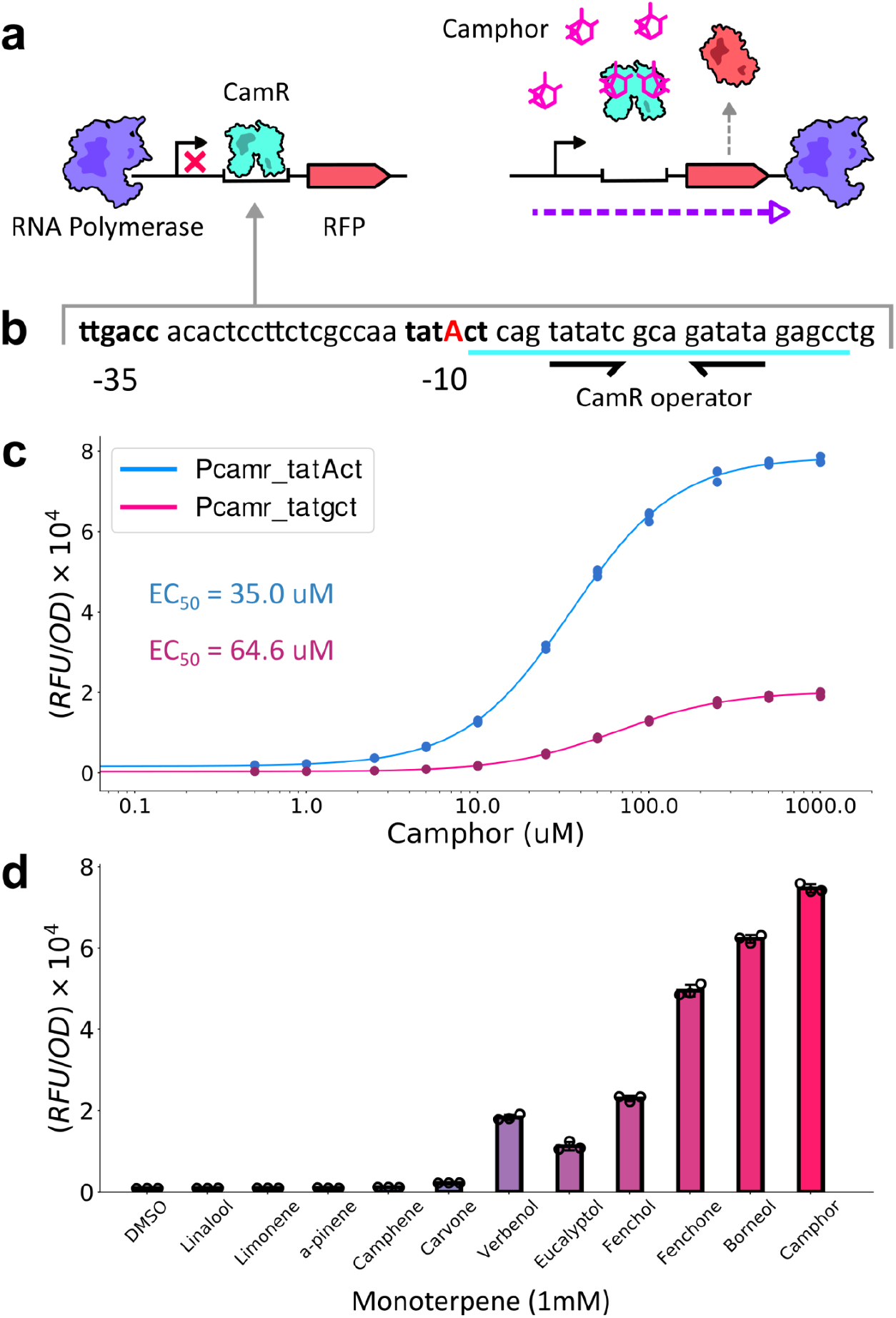
Design and characterization of the CamR biosensor system. (**a**) Schematic of the camphor-inducible biosensor circuit. (**b**) Sequence of the native P_camr_ promoter extracted from the native Pseudomonas putida plasmid. A G→A substitution was made in the -10 box (colored red) to increase transcription strength. Black arrows indicate the location of the inverted repeat. (**c**) Dose response function of the CamR biosensor system with the native promoter (tatgct) and the modified promoter (tatAct). EC_50_ values are listed and color coded (blue: tatAct, pink: tatgct) (**d**) Response of the CamR system to a panel of monoterpenes. All cultures were grown in the presence of 1% DMSO and fluorescence values are the averages of three biological replicates for both c and d.

A potential key enabler is the use of directed evolution to reconfigure the binding specificity of natural biosensors for new ligands^9,10,11^. Even so, directed evolution methods can be time-consuming, and often yield modest changes to effector responsiveness. We have now addressed the discovery gap by using a novel high-throughput selection and screening method (Seamless Enrichment of Ligand-Inducible Sensors, SELIS) to produce generalist biosensors that can be used across many different bicyclic monoterpenes. A camphor-responsive P_camr_/CamR system from *Pseudomonas putida* was repurposed to function in *Escherichia coli* and subsequent promoter optimization increased the system’s signal-to-noise ratio to 150-fold. SELIS was then used to evolve CamR towards four non-cognate monoterpenes, yielding rapid convergence on a series of broadly useful generalist biosensors.

## RESULTS

### Engineering a camphor-responsive biosensor and circuit

To identify a generalist monoterpene biosensor, we started from the camphor responsive CamR repressor from *P. putida*. This transcription factor has been thoroughly characterized *in vitro* [**9**], and camphor is structurally similar to other monoterpenes that have been produced microbially, including borneol^12^, fenchol^13^, eucalyptol^14^, and camphene^13^. The P_camr_ promoter was extracted from the natural *P. putida* CamR-bearing plasmid (GenBank: D14680.1), and included a portion protected by CamR in a DNase footprinting assay^15^ upstream from the self-cleaving ribozyme PlmJ^16^. To test the function of the heterologous P_camr_/CamR system in *E. coli*, CamR was constitutively expressed on one plasmid (pCamR) in the presence of a co-transformed plasmid bearing the P_camr_ promoter upstream of the mScarlet-i gene^17^ (pPcamr-RFP). The native P_camr_ promoter proved to be not very active in *E. coli* (**Figures 1b, 1c)**. We hypothesized that the -10 region of the promoter was divergent from the *E. coli* consensus ^11^, and introduced a single G→A base substitution in the -10 region, which greatly improved activity and the EC_50_ value (**Figures 1b, 1c**). Interestingly, while past attempts to repurpose the P_camr_/CamR system for camphor-inducible gene expression in *E. coli* have failed^18^, possibly due to an inhibitory effect of the native 5’-noncoding region of the native CamR mRNA transcript and insufficient transcription and translation strength, our largely synthetic design produced a reliable and robust dose-dependent signal.

The inherent response of CamR to different monoterpenes was analyzed. As expected, CamR was found to be highly responsive to its canonical effector, camphor, displaying an EC_50_ of 36.3 μM and a ∼40-fold signal-to-noise ratio (**Figure 1c**). Responses were also observed to a variety of bicyclic monoterpenes that contained alcohol, ketone, or ether functional groups (such as fenchol, fenchone, and eucalyptol), but not to other bicyclic, monocyclic, or acyclic monoterpenes (such as camphene, limonene, or linalool; **Figure 1d**).

We further explored the specificities of six different CamR homologs from various *Pseudomonas* species with the aim of identifying the best biosensor for subsequent evolution. The pCamR plasmid was accordingly modified to replace CamR with one of these six different homologs and was then co-transformed with the Pcamr-RFP plasmid prior for monoterpene screening, as described above. This screening effort revealed two other homologs that had a ligand profile similar to the *P. putida* CamR (mybCamR and thl2CamR), albeit with lower induction, and one homolog (tcuCamR) that was somewhat more specific for borneol and camphor **(Supplementary Figure 1**). Three homologs divergent from the *P. putida* CamR (CamV, my2CamR, bazCamR) generally displayed a poor response to all of the tested monoterpenes, suggesting that they might bind to an alternative operator sequence or have evolved to bind to other, currently unknown effector molecules.

### Optimization of reporter circuit performance

Given that the *P. putida* CamR functioned as a semi-specific biosensor, and that no other candidates appeared to be better generalists, we continued towards circuit optimization with this biosensor. To improve the performance of the circuit for resolving differences between and improvements with new, relatively inactive effector molecules, the dynamic range of the reporter system was systematically optimized via three separate components: the promoter and RBS adjacent to the reporter RFP, and the CamR operator sequence (**Supplementary Table 1**). Since strong promoters can produce a high background signal, while weak promoters result in a low induced signal (as was previously the case for the unmodified P_camr_ promoter), fine-tuning promoter strength can significantly improve the dynamic range of biosensor systems^19^. Eight synthetic promoters and five synthetic RBSs that each spanned a range of expression strengths about three orders of magnitude were introduced upstream of RFP. Multiple operator sequences were also considered during the optimization process, including the native operator (WT) and the inverted native operator (INV). In addition, since higher operator symmetry has previously been demonstrated to reduce background signal, possibly by increasing the strength of the regulator-DNA interaction^20^, two operator variants with greater symmetry than the native operator -- one located upstream of a CamR homolog (WP_145928353.1) in *Pseudomonas sp. TCU-HL1* (V2), and the other being our own synthetic design (V3) -- were introduced.

The medium-high strength P500 promoter produced the highest signal-to-noise ratio when cells were induced with 1 mM of camphor (**Figure 2b**), the synthetic operator sequence with the greatest symmetry (V3) surpassed all other tested operator sequences (largely by reducing background signal; **Figure 2c**), and the two strongest RBS sequences outperformed weaker counterparts (**Figure 2d**). Overall, after systematic optimization of the promoter, RBS, and operator, the best performing reporter circuit produced a 150-fold signal-to-noise ratio, ∼3.7-fold higher than the original construct (**Figure 2e**). The fact that a largely designed and synthetic circuit was functional in a heterologous system (*E. coli*) and outperformed the native system, primarily by reducing the background signal, reinforces the notion that synthetic parts may generally have greater orthogonality to the system as a whole, and hence greater predictability^21^.

**Figure 2.**
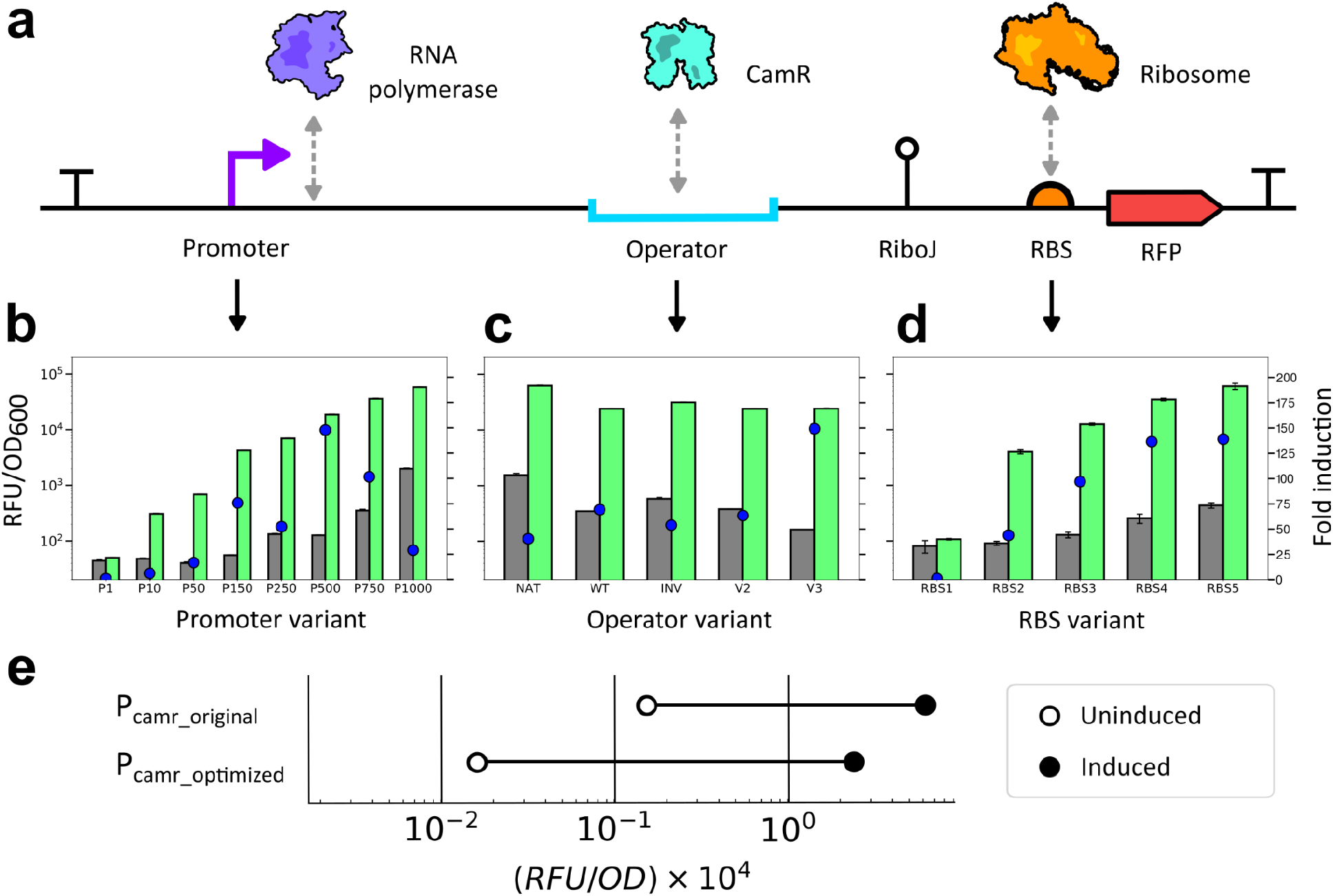
Systematic optimization of the Pcamr-RFP reporter. (**a**) Schematic of separate components of the P_camr_ reporter plasmid. (**b**) Fold induction of P_camr_ promoter variants. Grey bars and green bars represent the fluorescent signal of uninduced cells and cells induced with 1 mM camphor, respectively. Blue dots represent the fold induction of each variant in response to 1 mM camphor (**c**) Fold induction of P_camr_ operator variants. Same color scheme is used as for b. (**d**) Fold induction of P_camr_ RBS variants. Same color scheme is used as for b. (**e**) Fold induction of original P_camr_ promoter compared to the optimal P_camr_ promoter (P500, V3, RBS5) when induced with 1 mM of camphor.

### Evolution of CamR to encompass additional monoterpenes

While CamR was responsive to a number of bicyclic monoterpenes, further expanding its effector specificity would be greatly enabling for serving as a unitary biosensor for monitoring the metabolic engineering of strains to produce terpene products. We therefore leveraged a previously developed directed evolution approach, Seamless Enrichment of Ligand-Inducible Sensors (SELIS^22^, (**Supplementary Figure 2**), to improve the responsivity of CamR towards the terpene synthase products borneol, fenchol, eucalyptol, and camphene, all of which have been biosynthesized in *E. coli* previously^12,13,14^. SELIS relies on a positive screen for ligand-dependent regulator function via GFP production, and a conjoined negative selection in which transcriptional repression in the absence of ligand yields antibiotic resistance. Between the positive screen and negative selection, SELIS can quickly deconvolute libraries with over 10^5^ members in under a week and provides substantial genotype and phenotype data that allows optimally performing biosensors to be chosen by the researcher from amongst a range of successful candidates. CamR was introduced into the SELIS pipeline by placing the previously identified P_camr_ promoter construct upstream of a sfGFP gene for the positive screen and having a separate P_camr_ promoter drive the expression of the Lambda cI repressor for the negative selection (**Supplementary Figure 2**).

Initial CamR libraries were generated by site-saturating four sets of three, predicted, ligand-proximal residues, resulting in 32,000 unique protein mutants. When these largely failed to yield CamR variants responsive to the target monoterpenes, subsequent libraries were generated via error-prone mutagenesis of the entire CamR coding sequence, introducing an average of two mutations per gene. *E. coli* co-transformed with CamR libraries and the pSELIScamr plasmid were grown in the presence of zeocin (negative selection) and subsequently plated on solid media containing 1.0 % DMSO and either borneol, fenchol, eucalyptol, or camphene (positive screen; **Figure 3a**). Highly fluorescent clones were isolated, grown in the presence and absence of the target monoterpene, and ten clones with the high signal-to-noise ratios for each different effector were sequenced, a total of 40 CamR variants (**Supplementary Table 2**). It should be noted that since the fluorescence of clonal isolates is measured both with and without the effector during screening virtually all false positives are eliminated.

**Figure 3.**
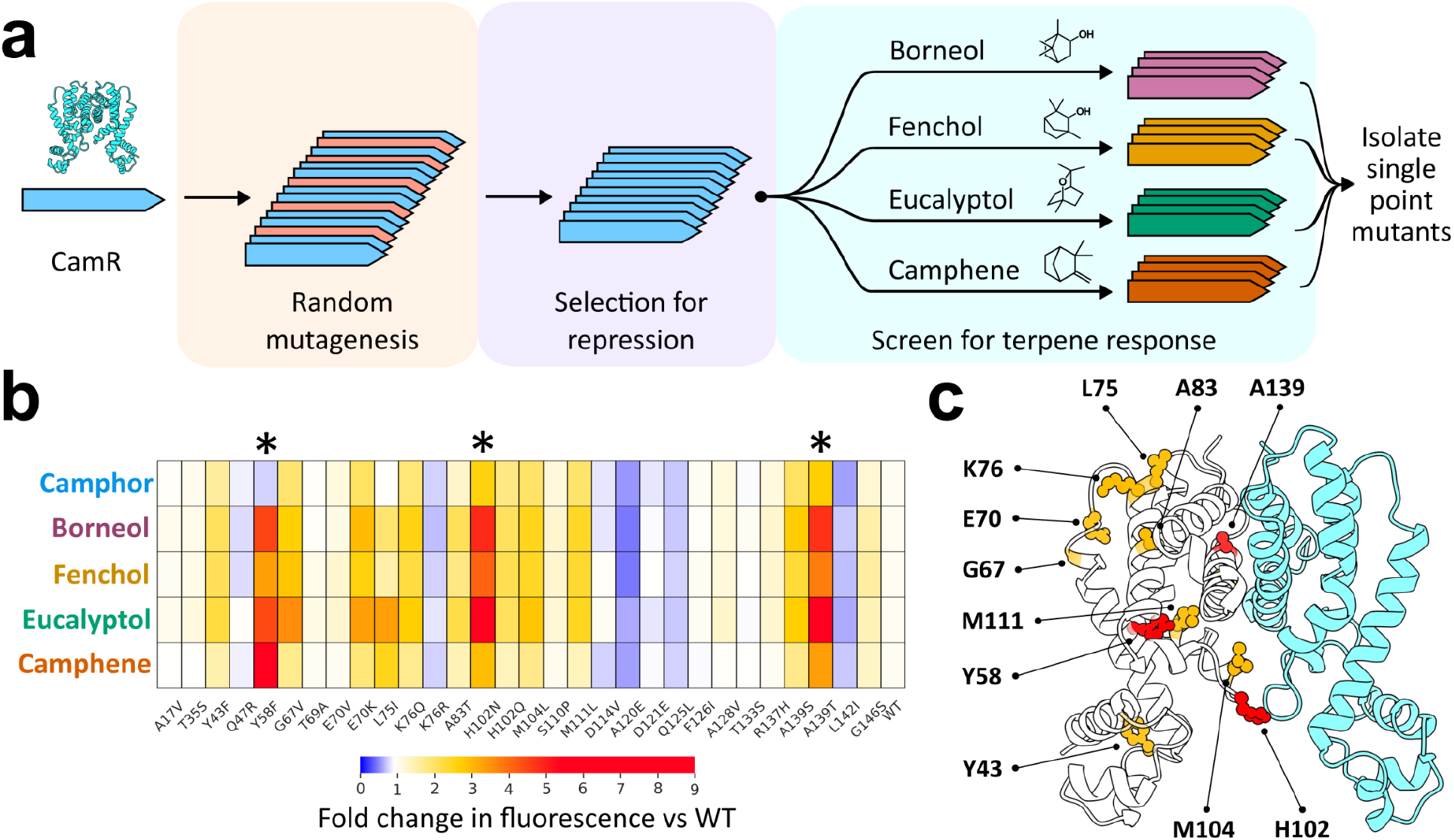
Evolution of CamR towards alternative monoterpenes produces generalists (**a**) Workflow for CamR evolution towards alternative monoterpenes. (**b**) Response of all recovered CamR single mutation variants to camphor, borneol, fenchol, eucalyptol, and camphene compared to the WT CamR protein. All cultures were grown in the presence of 1% DMSO and fluorescence values are the averages of three biological replicates. Asterisks indicate variants that produced a four-fold or greater change in fluorescent response. (**c**) Homology structure of CamR with single mutations recovered from evolution labelled on one dimer (left). Residues colored red correspond to mutants that increase the fold change in fluorescence four-fold or more, whereas residues colored orange correspond to mutants that increase the response by two to four-fold.

A large fraction of the 40 improved variants shared common substitutions. A Y58F substitution occurred in 25/40 of the recovered clones, while an A139T substitution appeared in 10/40 of the clones. Other shared substitutions, such as M111L, A83T, and G67V, were isolated in multiple screens for different monoterpenes (**Supplementary Table 1**).

Given that a majority (22/40) of the clones had only single substitutions, and that 34/40 clones had either the A139T or Y58F substitution, we hypothesized that we could best understand the major contributions to responsivity and specificity on a substitution-by-substitution basis. We therefore cloned all 30 unique single amino acid substitutions that were recovered from CamR evolution and screened them for activity against camphor, borneol, fenchol, eucalyptol, and camphene (**Figure 3b, Supplementary Figure 3**). As might have been expected based on the selection data alone, the three substitutions Y58F, H102N, and A139T independently were found to increase the response to at least one terpene by 6- to 7.5-fold. Ten other substitutions increased responsivity to at least one terpene by 2- to 3-fold, five increased the response by 20-50%, and 12 variants were found to not significantly increase the response to any terpenes. Interestingly, virtually every substitution that increased the response of CamR for one monoterpene also increased its response for all other tested monoterpenes to varying degrees. The two exceptions were Y58F and L75I, which improved responsiveness towards all terpenes except camphor.

To further understand the potential contributions of the individual amino acid changes, we mapped all 11 of the most productive substitutions onto a homology model of CamR (**Figure 3c**). With the exception of Y43 and A139 -- which are located in the DNA-binding and dimerization domains, respectively -- all nine other positions were located in the ligand binding domain. The predominant Y58F (and less fixed M111L) likely faces inward towards the presumed ligand binding cavity and may interact with the ligand directly (**Supplementary Figure 4**). The other mutated positions within the ligand binding domain are likely facing out towards the solvent, and thus their contributions towards altering terpene response are more difficult to rationalize.

While we had initially searched for a broadly useful generalist, the apparent lack of any variants that showed greater specificity for any of the monoterpenes was surprising, especially given previous efforts with SELIS to evolve a similar biosensor (RamR) to recognize benzylisoquinoline alkaloids. Since we and others have previously observed that the substrate- and effector specificities of evolving proteins often go through generalist intermediates prior to re-specialization^23,24^, we carried out two additional rounds of SELIS to see if we could increase responsiveness to fenchol **(Figure 4a**). While these additional generations were successively more responsive to fenchol, they also displayed an increased background signal (**Figure 4b**; to assist with analysis, CamR variants were expressed auto-inductively using the P_camr_ promoter during characterization (**Supplementary Figure 5**)). Interestingly, even as we attempted to force specialization on fenchol, the improved variants accumulated substitutions that had previously been observed in the CamR variants evolved for borneol, eucalyptol, and camphene, such as Y58F, H102N, and M104L (**Supplementary Figure 6** and **Supplementary Table 3**). Indeed, screening each final CamR generation against camphor, borneol, fenchol, eucalyptol, and camphene revealed that the later, fenchol-targeted generations had nonetheless become more broadly activated by all tested terpenes (**Figure 4c, Supplementary Figure 7**). Beyond these five monoterpenes, the final CamR generation, FEN3, was observed to respond more evenly to a wider range of bicyclic monoterpenes (**Figure 4d**).

**Figure 4.**
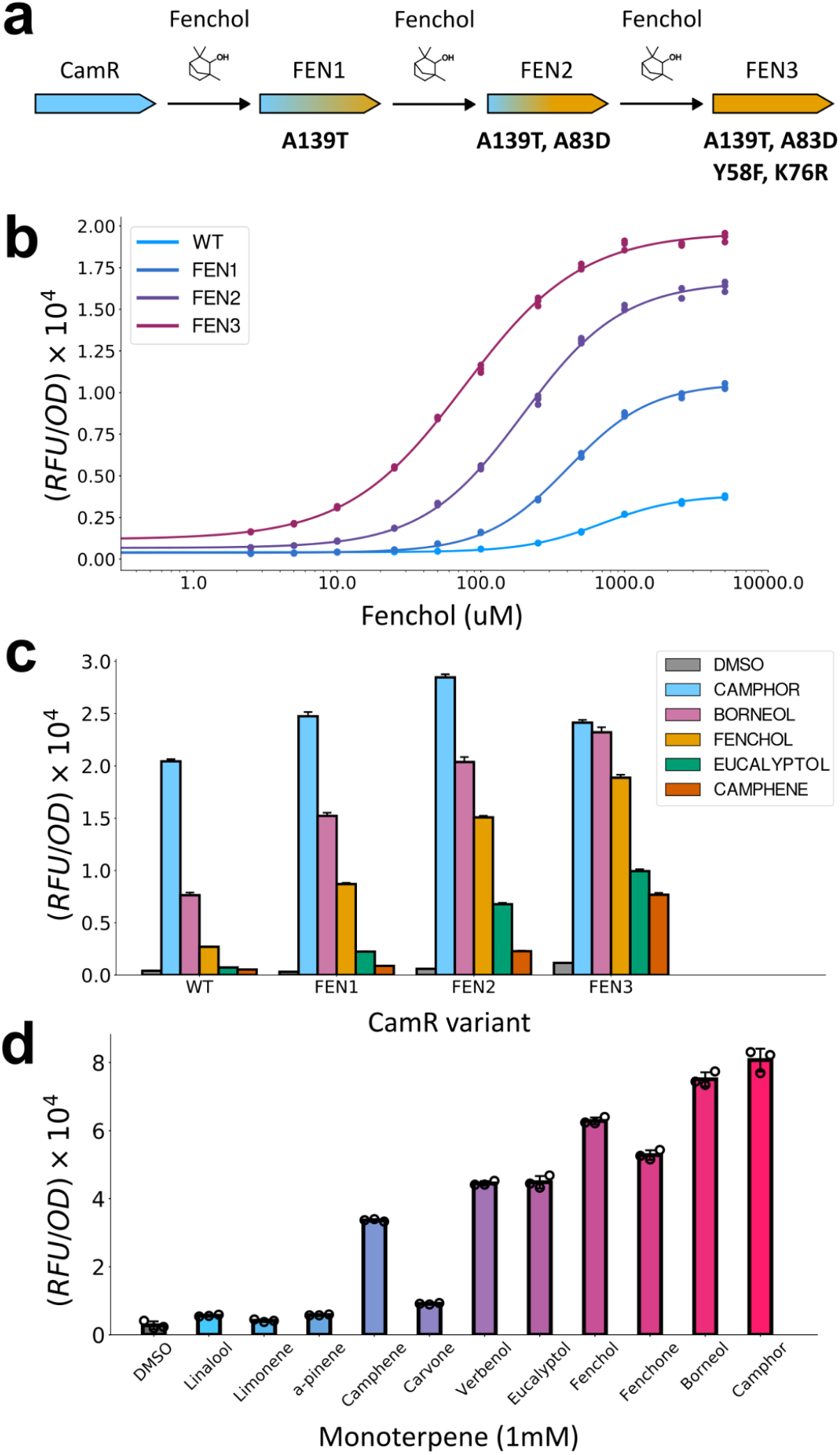
Evolution of CamR towards fenchol yields generalist monoterpene sensors (**a**) Workflow for multi-generation evolution of CamR towards fenchol. The genotypes of each variant are indicated in bold lettering. (**b**) Dose response function for each CamR generation with fenchol. (**c**) Response of each CamR generation to 1 mM of camphor, borneol, fenchol, eucalyptol, and camphene. (**d**) Fluorescent response of FEN3 to a panel of monoterpenes. All cultures were grown in the presence of 1% DMSO and fluorescence values are the averages of three biological replicates for panels **b, c**, and **d**.

## DISCUSSION

Genetic biosensors are gaining traction as high-throughput tools for otherwise difficult chemical analyses, such as distinguishing between compounds with identical exact masses and/or lacking a chromophore. However, biosensors often require custom engineering for target analytes, and even with great improvements in bioinformatic mining^4^ it is difficult to find corresponding biosensors. Therefore, we have developed a method (SELIS) that can be broadly used to identify biosensors, and herein we develop a genetic biosensor that is responsive to a range of bicyclic monoterpenes used in the flavor, fragrance, cosmetic, and pharmaceutical industries, which are traditionally limited to analysis using low-throughput gas chromatography-mass spectrometry. Indeed, microbes have already been engineered to produce many of the compounds recognized by our CamR sensors, such as borneol^12^, fenchol^13^, eucalyptol^14^, and camphene^13^, and our biosensors can now be used for metabolic engineering of such pathways, including identifying new monoterpene synthases^25^, improving strain health (since many monoterpenes are known to be toxic to microbes^26^), and balancing expression levels to reduce a heterologous pathway’s metabolic load^27^.

While we wanted to make one, broad-ranging sensor for the variety of monoterpene pathways and applications that are even now being developed, we were surprised to find that only generalists emerged from our selections, despite the application of both counterselection and additional rounds that attempted to ‘push’ a line towards fenchol specificity. These results stand in contrast to previous results with benzylisoquinoline alkaloids (BIAs), where we were able to find highly specific sensors for five different compounds starting from a similar TetR family member, RamR. The seeming intransigence of CamR to change, rather than broaden, effector specificity could be due to several factors. First, it is likely that different TetR family members are differentially evolvable. It is entirely possible that CamR has a narrow evolutionary landscape that is steeped in many, nearby generalist variants; this has previously been observed during the evolution of other transcriptional regulators^23,24^. Second, the ligand classes we have so far examined are in fact quite different, with BIAs occupying a much larger physical space than the monoterpenes explored herein (**Supplementary Figure 8**). Thus, there may just be more ‘handles’ for the effector binding site to recognize and distinguish for the BIAs, than for the monoterpenes. In this regard, it is interesting to note that the closest thing to a specialist that has appeared in nature or directed evolution is camphor-mediated activation of CamR itself, possibly because of the unique carbonyl on camphor. Third, the failure to find new specialists may be a deficiency in SELIS, in which the negative selection step is too stringent, and it is difficult for the protein to find an evolutionary path to a new specificity without going through intermediates that have high background in the absence of effector. This would reflect difficulties that have previously been observed in other coupled negative / positive selection schemes, such as those with tRNA synthetases^28^.

Irrespective of the mechanism, what is somewhat remarkable is that we are able to directly probe the evolvability of CamR via the straightforward directed evolution experiments that constitute SELIS, and hence draw out these hypotheses. Into the future, the evolutionary trajectories of CamR reported herein can be compared with other transcription factors to directly examine even more detailed hypotheses. For example, it has been suggested that protein evolvability correlates with both conformational plasticity within active site residues and rigid stability in core scaffold residues^29^. It is therefore possible that the ligand binding pocket of CamR is trapped in a rigid conformation and that mutations that occurred throughout our evolution experiments introduce flexibility and enable the exploration of a wider diversity of conformational states. The examination of the structures of intermediates and evolved variants can begin to address these questions.

Finally, the ability to quickly probe the evolutionary landscape of a target biosensor using SELIS will likely play an increasingly important role in guiding future sensor engineering efforts. Phylogenetic data combined with high-throughput experimentation can be leveraged to predict the capacity of transcription factor scaffolds to evolve for different ligand chemistries, creating a palette of the most useful starting points for biosensor development.

## METHODS

### Strains, plasmids, and media

*Escherichia coli* DH10B (New England BioLabs, Ipswich, MA, USA) was used for all routine cloning and directed evolution. All biosensor systems were characterized in *E. coli* DH10B. LB-Miller (LB) media (BD, Franklin Lakes, NJ, USA) was used for routine cloning, fluorescence assays, directed evolution, and orthogonality assays unless specifically noted. LB + 1.5% agar (BD, Franklin Lakes, NJ, USA) plates were used for routine cloning and directed evolution. The plasmids described in this work were constructed using Gibson assembly, golden gate assembly, and standard molecular biology techniques. Synthetic genes, obtained as gBlocks, and primers were purchased from IDT. Relevant plasmid sequences are provided in **Supplementary Table 4** and are available by request from the corresponding authors.

### Monoterpenes

Cells were induced with the following chemicals: Camphor (Tokyo Chemical Industry, CAT#: C0011); borneol (Tokyo Chemical Industry, CAT#: B0525); fenchone (Tokyo Chemical Industry, CAT#: F0164); fenchol (Alfa Aesar, CAT#: L03211); eucalyptol (Acros Organics, CAT#: 110340050); verbenol (Sigma Aldrich, CAT#: 247065-5G); camphene (Tokyo Chemical Industry, CAT#: C0009); carvone (Acros Organics, CAT#: 154590050); alpha-pinene (Acros Organics, CAT#: 131270050); limonene (Tokyo Chemical Industry, CAT#: L0132); linalool (Combi-Blocks, CAT#: QH-3254). All monoterpenes were flushed with argon and stored at -20°C after opening. It should be noted that the commercially available camphene used in this study is >78% pure and contains as much as 20% tricyclene.

### Chemical transformation

For routine transformations, strains were made competent for chemical transformation. 5 mL of an overnight culture of DH10B cells were subcultured into 500 mL of LB media and grows at 37°C, 250 r.p.m. for 3 h. Cultures were centrifuged (3,500 g, 4 °C, 10 min), and pellets were washed in 70 mL of chemical competence buffer (10% glycerol, 100 mM CaCl2) and centrifuged again (3,500 g, 4°C, 10 min). The resulting pellets were resuspended in 20 mL of chemical competence buffer. After 30 minutes on ice, cells were divided into 250 μL aliquots and flash frozen in liquid nitrogen. Competent cells were stored at −80 °C until use.

### Promoter design

The original Pcamr promoter (see supplementary figure 1) was derived from the literature [**ref**]. For modified Pcamr promoters, either the promoter, operator, or RBS sequence was swapped using golden gate assembly (**Supplementary Table 1**). The RiboJ insulator was used in all promoter designs to insulate the variable operator context and increase the overall fluorescent signal ^16^. Promoter sequences were based on the J23100 Andersen promoter and either the upstream activating sequence, the -35 region (TTGACA), or the -10 region (TATAAT) were modified to afford variable expression strengths. RBS sequences were designed using the RBS calculator ^30^. A terminator was placed immediately upstream from the promoter sequence to insulate the reporter output from upstream transcriptional activity (see **Supplementary Table 4**).

### Biosensor response measurement

For performing a biosensor response assay, the pCamR and pPcamr-RFP plasmids were co-transformed into DH10B cells and plated on an LB agar plate with appropriate antibiotics. Three separate colonies were picked for each transformation and were grown overnight. The following day, 20 μL of each culture was then used to inoculate 12 separate wells within a 2 mL 96-deep-well plate (Corning, Product #: P-DW-20-C-S) sealed with an AeraSeal film (Excel Scientific, Victorville, CA, USA) containing 900 μL of LB media. After two hours of growth at 37 °C cultures were induced with 100 μL of LB media containing either 10 μL of just DMSO or the target monoterpene dissolved in 10 μL of DMSO. Cultures were grown for an additional 4 hours at 37 °C, 250 r.p.m and subsequently centrifuged (3,500 g, 4°C, 10 min). Supernatant was removed and cell pellets were resuspended in 1 mL of PBS (137 mM NaCl, 2.7 mM KCl, 10 mM Na2HPO4, 1.8 mM KH2PO4. pH 7.4). One hundred μL of the cell resuspension for each condition was transferred to a 96 well microtiter plate (Corning, Product #: 3904), from which the fluorescence (Ex: 566 nm, Em: 596 nm) and absorbance (600 nm) was measured using the Tecan Infinite M1000 plate reader.

### CamR library construction

Random mutagenesis libraries were created by amplifying the coding region of the CamR gene with error-prone polymerase chain reaction. To perform this mutagenesis protocol, 50 ng of the template gene was mixed with 2 uL of 2 uM forward and reverse primers, 3 uL of Taq polymerase (New England BioLabs, Ipswich, MA, USA), 1 uM of 50 mM MnCl_2_, 41 uL of water, and 50 uL of a custom buffer (300 uL 10x Taq polymerase buffer, 16.5 uL 1 M MgCl_2_, 15 uL 10 mg/mL bovine serum albumin, 6 uL 100 mM dGTP, 11 uL 100 mM dATP, 12 uL 100 mM dCTP, 40.5 uL 100 mM dTTP, and 1099 uL water). The thermal cycling procedure is as follows: 94 °C for 30 seconds, 55C for 30 seconds, 72C for one minute, repeated for 25 cycles. Libraries were cloned into the pCamR plasmid using Gibson assembly and *E. coli* DH10B bearing pSELIScamr was transformed with the resulting library. Transformation efficiency always exceeded 10^6^ for each round of selection, indicating several fold coverage of the library. Transformed cells were grown in LB media overnight at 37°C in carbenicillin and chloramphenicol.

### Directed evolution and validation of CamR biosensors

Twenty μL of cell culture bearing the sensor library was seeded into 5 mL of fresh LB containing appropriate antibiotics and 100 μg/mL zeocin (Thermo Fisher. CAT#: R25001) and were grown at 37°C for seven hours. Following incubation, 0.5 μL of culture was diluted into 1 mL of LB media, from which 100 μL was further diluted into 900 μL of LB media. Three hundred μL of this mixture was then plated across three LB agar plates containing carbenicillin, chloramphenicol and the target monoterpene dissolved in DMSO. Plates were incubated overnight at 37 °C. The following day the brightest colonies were picked and grown overnight in 1 mL of LB media containing appropriate antibiotics within a 96-deep-well plate sealed with an AeraSeal film at 37°C. A glycerol stock of cells containing pSELIScamr and pCamR bearing the parental CamR variant was also inoculated in 5 mL of LB for overnight growth.

The following day, 20 μL of each culture was used to inoculate two separate wells within a new 96-deep-well plate containing 900 μL of LB media. Additionally, eight separate wells containing 1 mL of LB media were inoculated with 20 μL of the overnight culture expressing the parental CamR variant. A typical arrangement would have 44 unique clones on the top half of the plate, duplicates of those clones on the bottom half of the plate, and the right-most column occupied by cells harboring the parental CamR variant. After 2 hours of growth at 37°C the top half of the 96-well plate was induced with 100 μL of LB media containing 10 μL of DMSO whereas the bottom half of the plate was induced with 100 μL of LB media containing the target monoterpene dissolved in 10 μL of DMSO. The concentration of BIA used for induction is typically the same concentration used in the LB agar plate for screening during that particular round of evolution. Cultures were grown for an additional 4 hours at 37°C, 250 r.p.m and subsequently centrifuged (3,500 g, 4°C, 10 min). Supernatant was removed and cell pellets were resuspended in 1 mL of PBS. One hundred μL of the cell resuspension for each condition was transferred to a 96 well microtiter plate, from which the fluorescence (Ex: 485 nm, Em: 509 nm) and absorbance (600 nm) was measured using the Tecan Infinite M1000. Clones with the highest signal-to-noise ratio were then sequenced and subcloned into a fresh pCamR vector.

For single mutation analysis, individual mutations were separately introduced within the CamR coding region of the pCamR plasmid. These plasmids were co-transformed with Pcamr-RFP, plated on solid media containing appropriate antibiotics, and the following day three individual colonies from each transformation were grown overnight. The resulting cultures were then assayed, as described in “Biosensor response measurement”, using 1 mM of each monoterpene dissolved in 10 μL of DMSO.

For sensor variant dose response measurement, each CamR variant was first subcloned into a fresh pCamR plasmid backbone. The resulting plasmids were then transformed into DH10B cells bearing pPcamr-RFP and three individual colonies from each transformation were subsequently grown overnight. The resulting cultures were then assayed, as described in “Biosensor response measurement”, using seven to eleven different concentrations of the target monoterpene along a DMSO-only control. For further evolution, sensor variants that displayed a combination of a low background, a reduced EC_50_ for the target monoterpene, and a high signal-to-noise ratio were used as templates for the next round of evolution.

### Homology model

The homology model of CamR was constructed using SWISS-MODEL (https://swissmodel.expasy.org/). 6AYI was used as a template for structure generation.

### Statistical analysis and reproducibility

All data in the manuscript are displayed as mean ± s.e.m. unless specifically indicated. Bar graphs, dose response functions, and orthogonality matrices were all plotted in Python 3.6.9 using matplotlib and seaborn. Dose response curves and EC_50_ values were estimated by fitting to the hill equation y = d + (a-d)*x^b^ / (c^b^ + x^b^) (where y = output signal, b = hill coefficient, x = ligand concentration, d = background signal, a = the maximum signal, and c = the EC_50_), with the scipy.optimize.curve_fit library in Python.

## Acknowledgements

Funding from DARPA Soils (HR00111920019), Welch (F-1654), and AFSOR - (FA9550-14-1-0089) is acknowledged.

## Author contributions

S.D. designed and performed all experiments. V.N. constructed and validated the promoter and RBS series. The manuscript was written by S.D. with support from A.D.E and H.A. S.D., A.D.E., and H.A. supervised all aspects of the study.

## Supplementary information

Additional experimental details, including circuit designs and biosensor genotype and phenotype data.

## References

1. Anuj Pathak. High-Throughput Screening: Technologies and Global Markets. BCC Research. PHM205A (2019).

2. Soares-Castro, P., Soares, F. & Santos, P. M. Current Advances in the Bacterial Toolbox for the Biotechnological Production of Monoterpene-Based Aroma Compounds. Molecules 26, 91 (2021).

3. Lin, J.-L., Wagner, J. M. & Alper, H. S. Enabling tools for high-throughput detection of metabolites: Metabolic engineering and directed evolution applications. Biotechnol. Adv. 35, 950–970 (2017).

4. Hanko, E. K. R. et al. A genome-wide approach for identification and characterisation of metabolite-inducible systems. Nat. Commun. 11, 1213 (2020).

5. Siu, Y., Fenno, J., Lindle, J. M. & Dunlop, M. J. Design and Selection of a Synthetic Feedback Loop for Optimizing Biofuel Tolerance. ACS Synth. Biol. 7, 16–23 (2018).

6. Phoenix, P. et al. Characterization of a new solvent-responsive gene locus in Pseudomonas putida F1 and its functionalization as a versatile biosensor. Environ. Microbiol. 5, 1309–1327 (2003).

7. Fujita, M., Aramaki, H., Horiuchi, T. & Amemura, A. Transcription of the cam operon and camR genes in Pseudomonas putida PpG1. J. Bacteriol. 175, 6953–6958 (1993).

8. Eaton, R. W. p-Cymene catabolic pathway in Pseudomonas putida F1: cloning and characterization of DNA encoding conversion of p-cymene to p-cumate. J. Bacteriol. 179, 3171–3180 (1997).

9. Yao, J. et al. Developing a highly efficient hydroxytyrosol whole-cell catalyst by de-bottlenecking rate-limiting steps. Nat. Commun. 11, 1515 (2020).

10. Snoek, T. et al. Evolution-guided engineering of small-molecule biosensors. Nucleic Acids Res. 48, e3–e3 (2020).

11. Xiong, D. et al. Improving key enzyme activity in phenylpropanoid pathway with a designed biosensor. Metab. Eng. 40, 115–123 (2017).

12. Lei, D. et al. Combining Metabolic and Monoterpene Synthase Engineering for de Novo Production of Monoterpene Alcohols in Escherichia coli. ACS Synth. Biol. 10, 1531–1544 (2021).

13. Leferink, N. G. H. et al. A ‘Plug and Play’ Platform for the Production of Diverse Monoterpene Hydrocarbon Scaffolds in Escherichia coli. ChemistrySelect 1, 1893–1896 (2016).

14. Mendez-Perez, D. et al. Production of jet fuel precursor monoterpenoids from engineered Escherichia coli. Biotechnol. Bioeng. 114, 1703–1712 (2017).

15. Aramaki, H., Sagara, Y., Hosoi, M. & Horiuchi, T. Evidence for autoregulation of camR, which encodes a repressor for the cytochrome P-450cam hydroxylase operon on the Pseudomonas putida CAM plasmid. J. Bacteriol. 175, 7828–7833 (1993).

16. Lou, C., Stanton, B., Chen, Y.-J., Munsky, B. & Voigt, C. A. Ribozyme-based insulator parts buffer synthetic circuits from genetic context. Nat. Biotechnol. 30, 1137–1142 (2012).

17. Bindels, D. S. et al. mScarlet: a bright monomeric red fluorescent protein for cellular imaging. Nat. Methods 14, 53–56 (2017).

18. Aramaki, H., Fujita, M., Sagara, Y., Amemura, A. & Horiuchi, T. Heterologous expression of the cytochrome P450cam hydroxylase operon and the repressor gene of Pseudomonas putida in Escherichia coli. FEMS Microbiol. Lett. 123, 49–54 (1994).

19. Chen, Y. et al. Tuning the dynamic range of bacterial promoters regulated by ligand-inducible transcription factors. Nat. Commun. 9, 64 (2018).

20. Ruegg, T. L. et al. Jungle Express is a versatile repressor system for tight transcriptional control. Nat. Commun. 9, 3617 (2018).

21. Temme, K., Zhao, D. & Voigt, C. A. Refactoring the nitrogen fixation gene cluster from Klebsiella oxytoca. Proc. Natl. Acad. Sci. 109, 7085–7090 (2012).

22. d’Oelsnitz, S. et al. Using structurally fungible biosensors to evolve improved alkaloid biosyntheses. (2021) doi:10.1101/2021.06.07.447399.

23. Collins, C. H., Arnold, F. H. & Leadbetter, J. R. Directed evolution of Vibrio fischeri LuxR for increased sensitivity to a broad spectrum of acyl-homoserine lactones. Mol. Microbiol. 55, 712–723 (2005).

24. Taylor, N. D. et al. Engineering an allosteric transcription factor to respond to new ligands. Nat. Methods 13, 177–183 (2016).

25. Leferink, N. G. H. et al. An automated pipeline for the screening of diverse monoterpene synthase libraries. Sci. Rep. 9, 11936 (2019).

26. Zhang, L. et al. Chassis and key enzymes engineering for monoterpenes production. Biotechnol. Adv. 35, 1022–1031 (2017).

27. Hartline, C. J., Schmitz, A. C., Han, Y. & Zhang, F. Dynamic control in metabolic engineering: Theories, tools, and applications. Metab. Eng. 63, 126–140 (2021).

28. Thyer, R. et al. Directed Evolution of an Improved Aminoacyl-tRNA Synthetase for Incorporation of L-3,4-Dihydroxyphenylalanine (L-DOPA). Angew. Chem. 133, 14937–14942 (2021).

29. Tóth-Petróczy, Á. & Tawfik, D. S. The robustness and innovability of protein folds. Curr. Opin. Struct. Biol. 26, 131–138 (2014).

30. Salis, H. M., Mirsky, E. A. & Voigt, C. A. Automated design of synthetic ribosome binding sites to control protein expression. Nat. Biotechnol. 27, 946–950 (2009).

