## Supplementary Information for "Evolving a generalist biosensor for bicyclic monoterpenes"

Supplementary Figures 1-8

Supplementary Tables 1-4

### SUPPLEMENTARY FIGURES

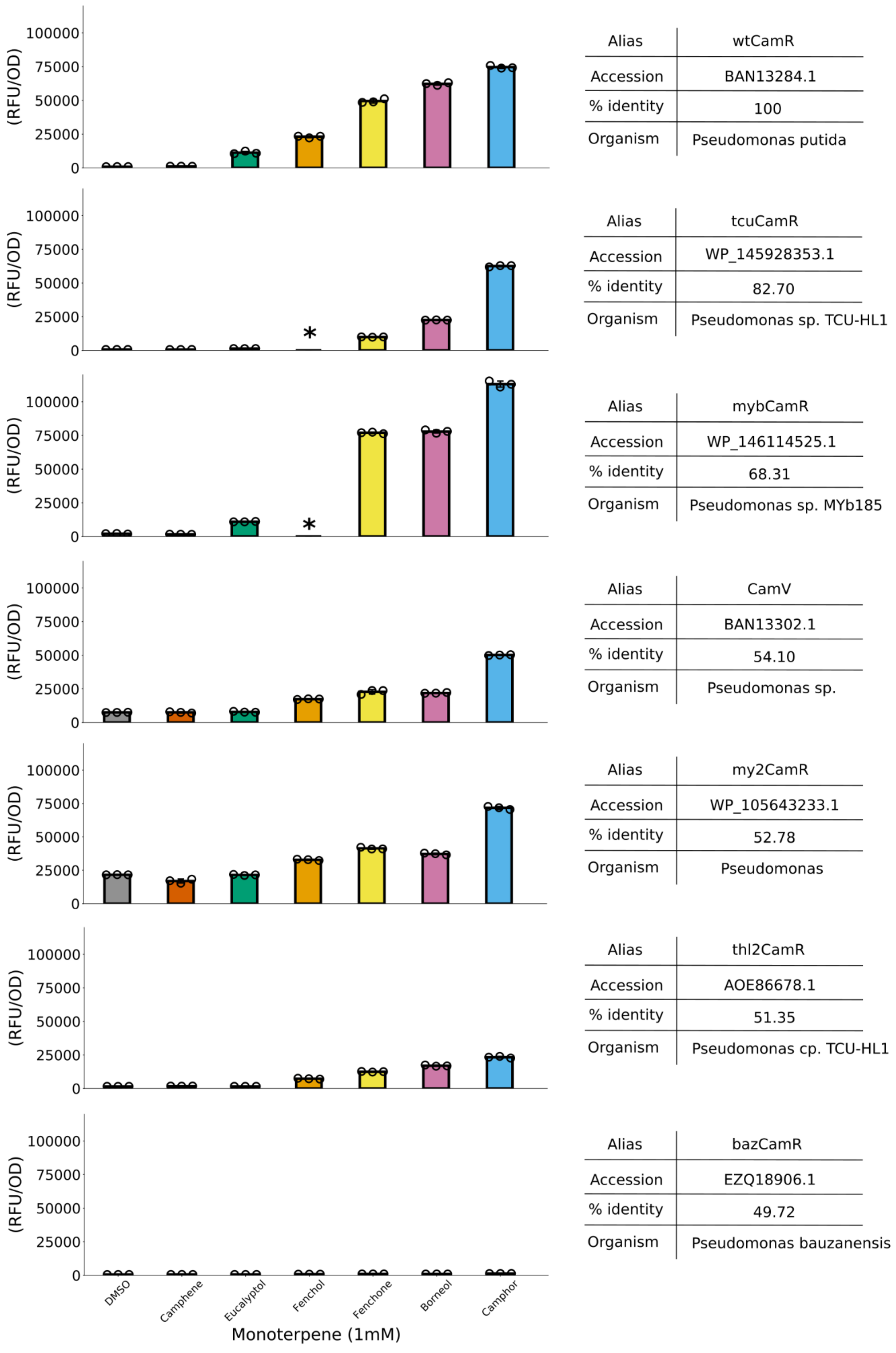


**Supplementary Figure 1.** Ligand specificity of CamR homologs.

Six CamR homologs from different pseudomonas species were screened for response to the bicyclic monoterpenes camphene (vermillion), eucalyptol (blue-green), fenchol (orange), fenchone (yellow), borneol (purple), and camphor (blue) and were compared to a solvent-only control (grey). Cells were induced to a final concentration of 1 mM of the target monoterpene and 1% DMSO. All assays were performed in biological triplicate. The alias, protein accession number, percent identity to wild-type CamR, and host organism are displayed to the right of the response data. Asterisks represent that data for that condition was not collected.

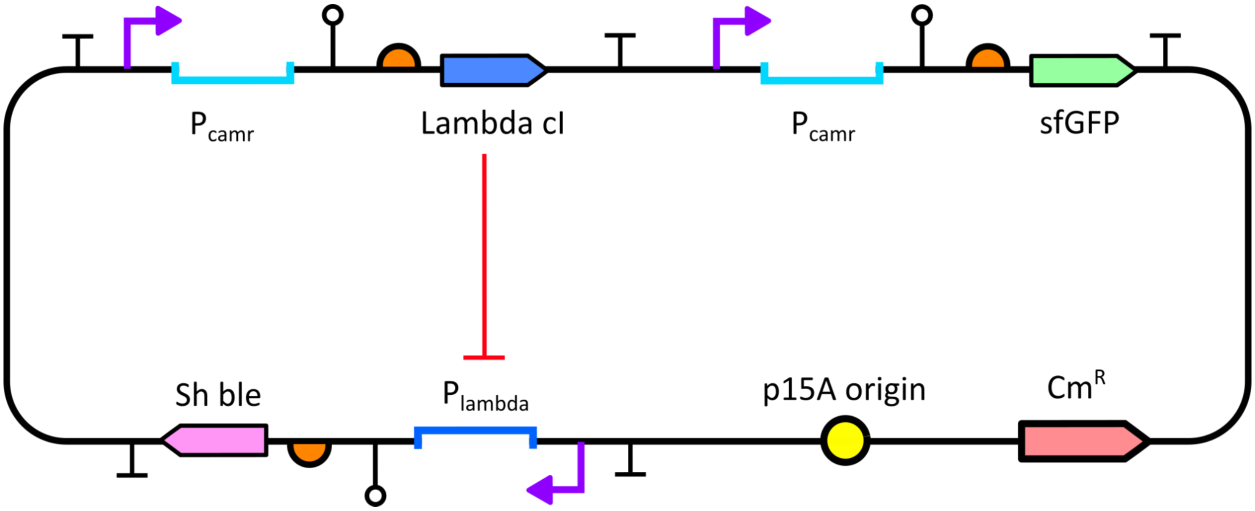


**Supplementary Figure 2.** Schematic of the pSELIScamr plasmid used for CamR evolution.

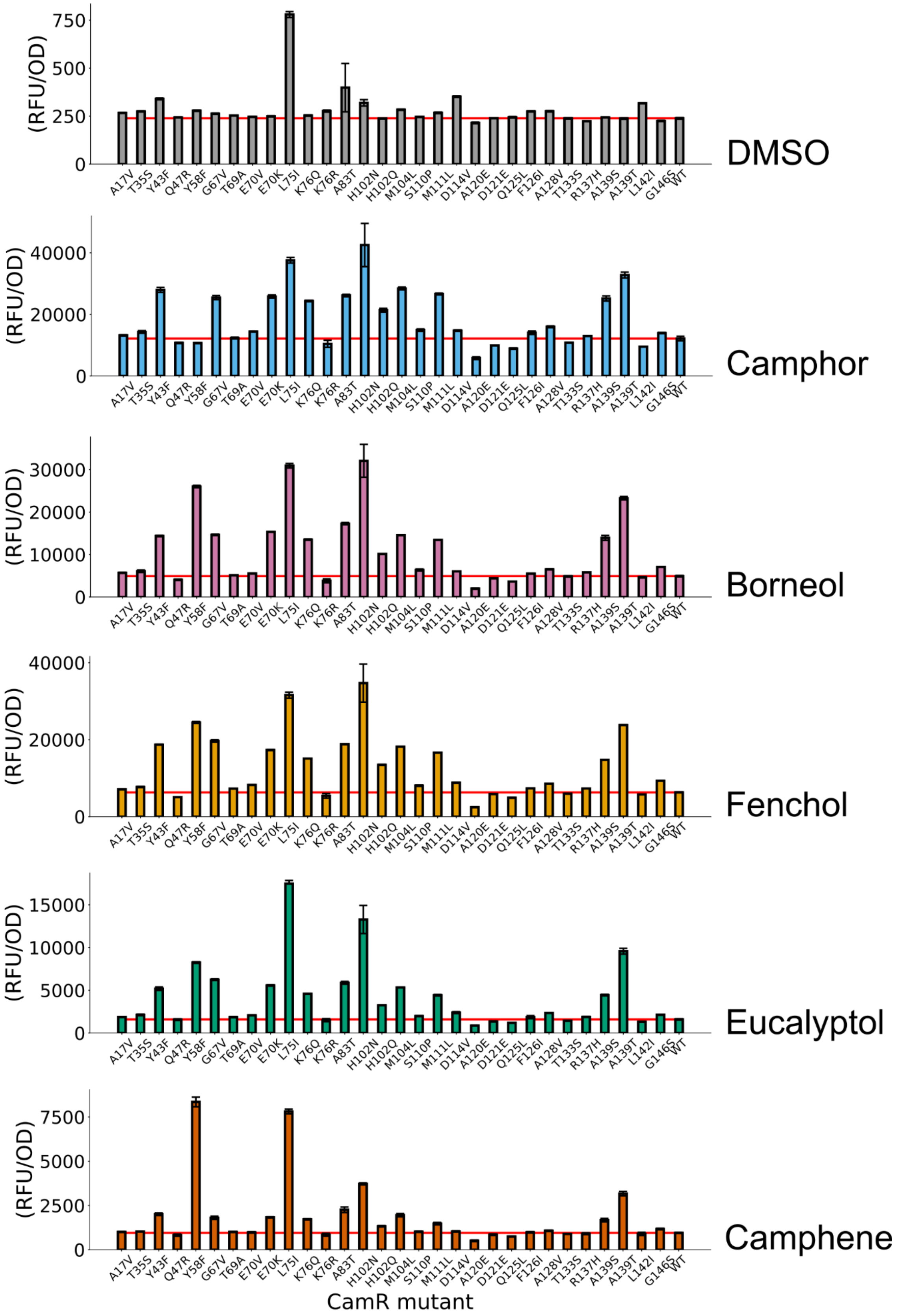


**Supplementary Figure 3.** Response of CamR single point mutants to all test monoterpenes. All measurements were performed in biological triplicate. Error bars represent the SE +/- the mean. The red horizontal line indicates the mean fluorescence of the natural template CamR sensor.

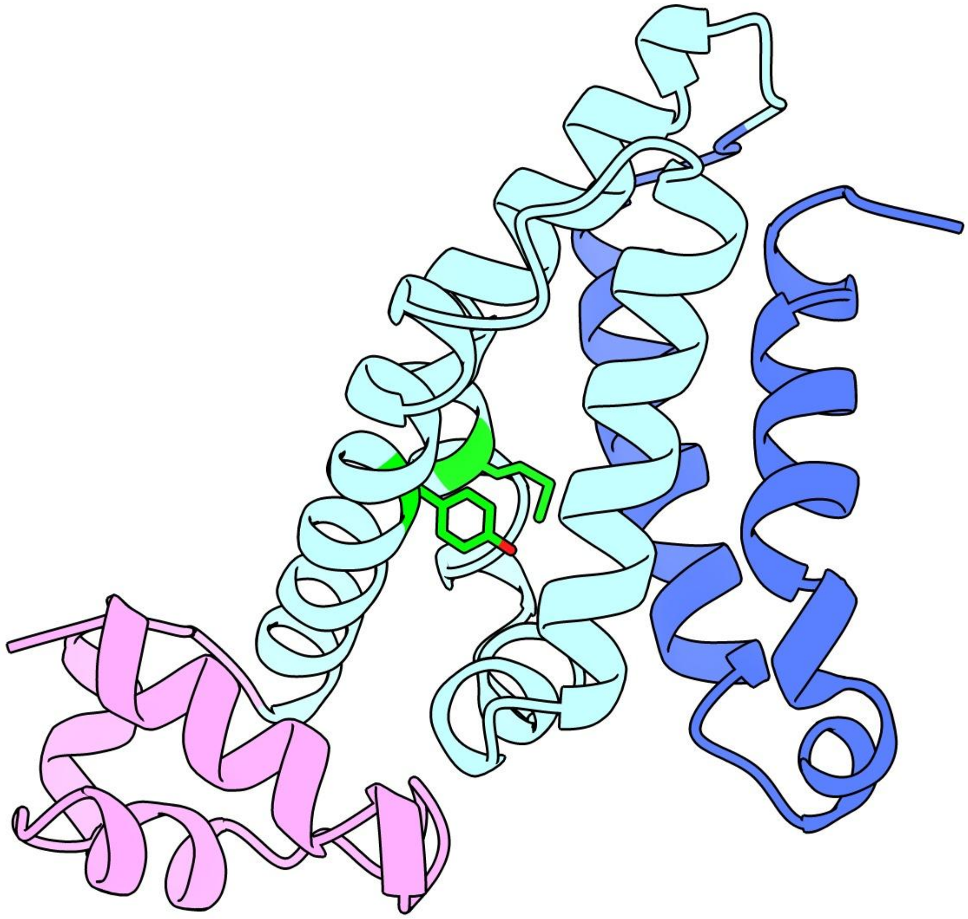

**Supplementary Figure 4.** Local environment of the Y58 and M111 residues.
The homology structure of one dimer of CamR. The DNA binding domain is shown in pink, ligand binding domain in cyan, dimerization domain in blue, and the Y58 and M111 residues in green.

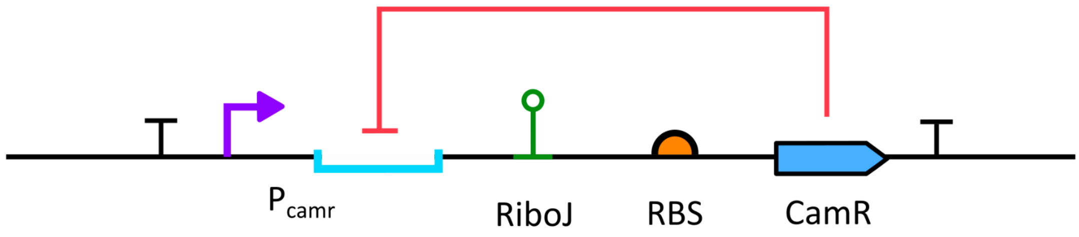


ggctgtcgacctttgaaaagttcgTTTACAgctagctcagtcctaggTACAATCTCACTATATCGCCGATATAGAGCCTactgagcacagctggctgagcacAGCTGTC
ACCGGATGTGCTTTCCGGTCTGATGAGTCCGTGAGGACGAAACAGCCTCTACAAATAATTTTGTTTAAactagTGAACCACGAGGCCTacatATG

**Supplementary Figure 5.** Schematic of the CamR autoinduction circuit and promoter sequence. Colors of each element are as follows: promoter sequence (purple), operator sequence (cyan), RiboJ insulator sequence (green), RBS sequence (orange), and CamR coding sequence (blue). The last three bases (ATG) represent the start of the coding sequence of CamR.

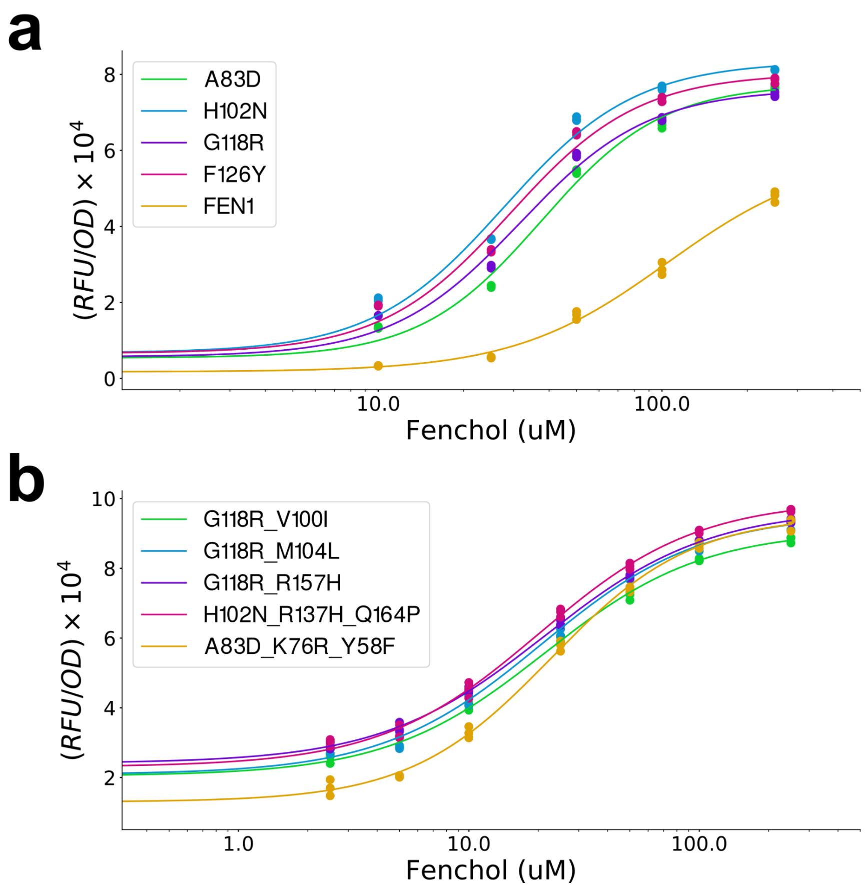


**Supplementary Figure 6.** Dose responses for all top performing CamR variants evolved for fenchol (**a**) Dose response functions of the top unique second generation CamR variants. All genotypes in the figure legend are new mutations that emerged in the context of the A139T mutant. (**b**) Dose response function of the top unique third generation CamR variants evolved for fenchol. All genotypes in the figure legend are new mutations that emerged in the context of the A139T, A83D mutant. All variants were subcloned into a new pCamR backbone prior to characterization with the $pP_{camr}$-RFP plasmid. All fluorescence measurements were performed in biological triplicate.

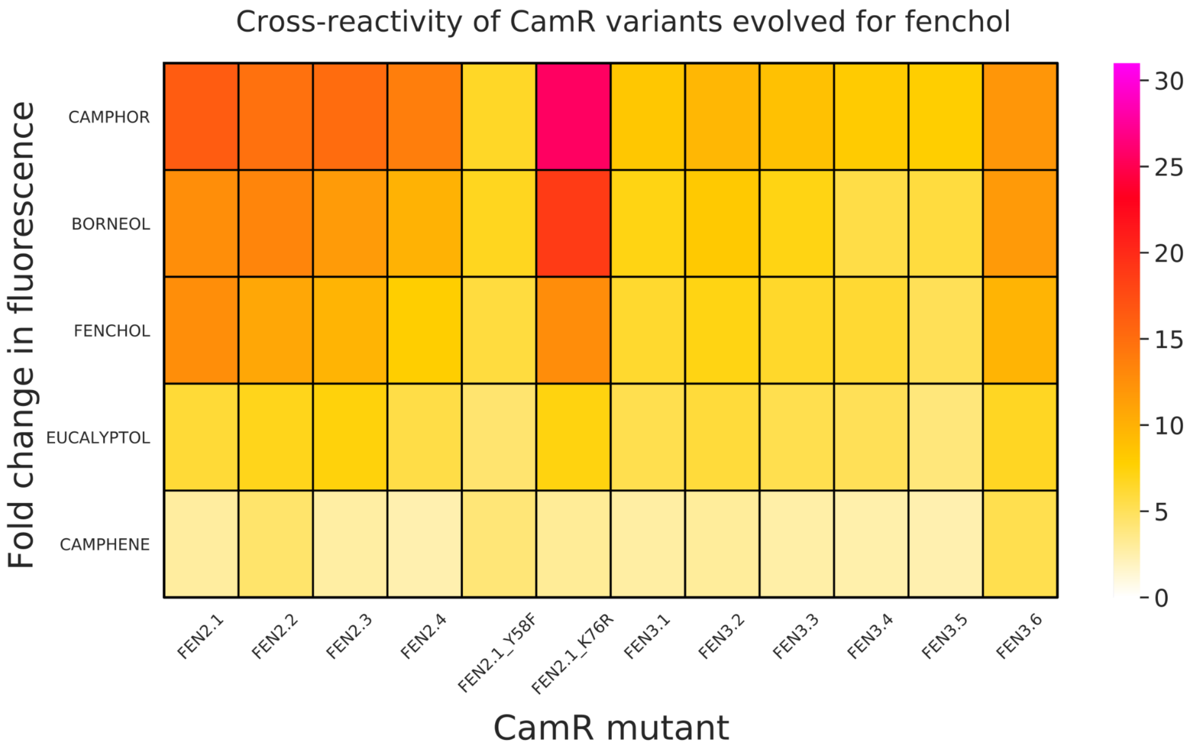


**Supplementary Figure 7.** Monoterpene specificity of all evolved CamR variants
All CamR variants were induced with 1mM of the monoterpene indicated. Fold change values are relative to a control condition induced with 10 uL of the solvent DMSO alone. The genotype of each variant can be found in Supplementary Table 3.

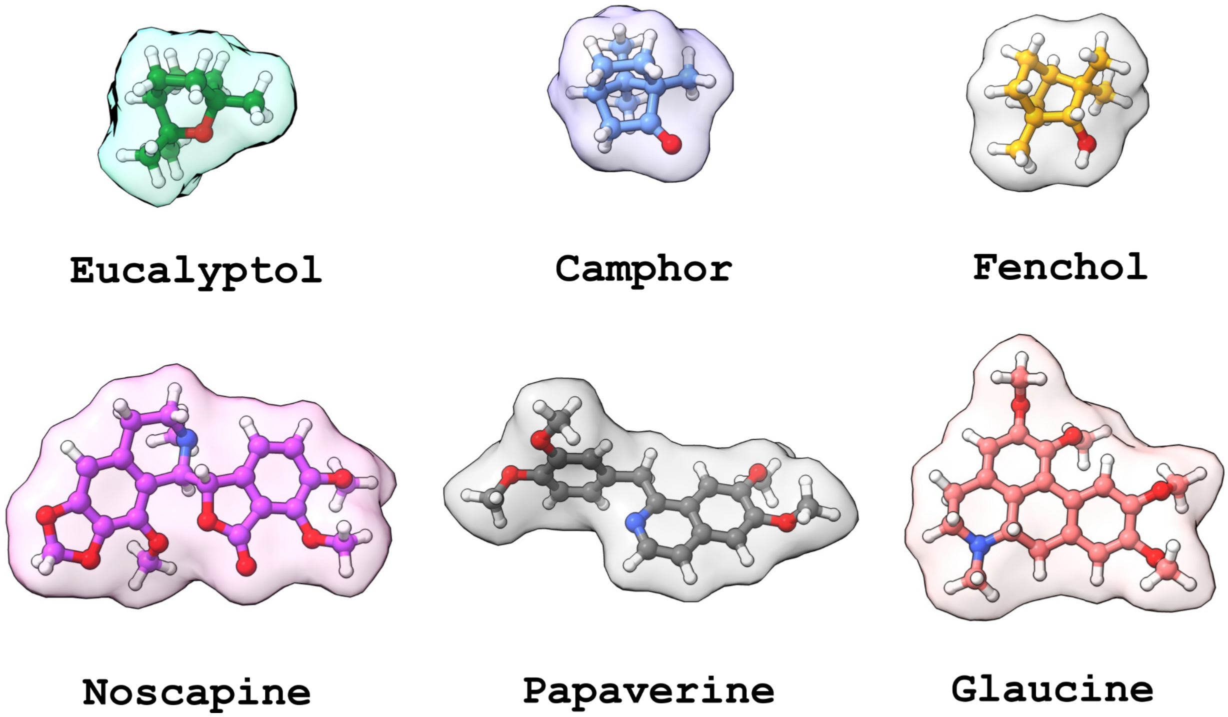


**Supplementary Figure 8**. 3D structures of representative monoterpenes compared to benzylisoquinoline alkaloids. Structures are displayed as ball-and-stick models using ChimeraX.

### SUPPLEMENTARY TABLES

#### Supplementary Table 1: Sequences of all Pcamr promoter variants

| Pcamr variant | Promoter |
|---|---|
| P1 | GGCTGTCGACCTTTGAAAAGTTCG**ATTACA**GCTAGCTCAGTCCTAGG**GACAAT**ctcactatatcgccgatatagagcctactgagcacagctggctgagcacagctgtcaccggatgtgctttccggtctgatgagtccgtgaggacgaaacagcctctacaaataattttgtttaaactagaagtaagcgaggtacacatATG |
| P10 | GGCTGTCGACCGGCGAAAAGTTCG**ATTACA**GCTAGCTCAGTCCTAGG**TACAAT**ctcactatatcgccgatatagagcctactgagcacagctggctgagcacagctgtcaccggatgtgctttccggtctgatgagtccgtgaggacgaaacagcctctacaaataattttgtttaaactagaagtaagcgaggtacacatATG |
| P50 | GGCTGTCGACCGGCGTGCCGTTCG**TTTACC**GCTAGCTCAGTCCTAGG**TACAAT**ctcactatatcgccgatatagagcctactgagcacagctggctgagcacagctgtcaccggatgtgctttccggtctgatgagtccgtgaggacgaaacagcctctacaaataattttgtttaaactagaagtaagcgaggtacacatATG |
| P150 | GGCTGTCGACCTTTGAAAAGTTCG**TTTACC**GCTAGCTCAGTCCTAGG**TACAAT**ctcactatatcgccgatatagagcctactgagcacagctggctgagcacagctgtcaccggatgtgctttccggtctgatgagtccgtgaggacgaaacagcctctacaaataattttgtttaaactagaagtaagcgaggtacacatATG |
| P250 | AAAATATATTTTTCAAAAGTATCG**TTTACC**GCTAGCTCAGTCCTAGG**TACAAT**ctcactatatcgccgatatagagcctactgagcacagctggctgagcacagctgtcaccggatgtgctttccggtctgatgagtccgtgaggacgaaacagcctctacaaataattttgtttaaactagaagtaagcgaggtacacatATG |
| P500 | GGCTGTCGACCTTTGAAAAGTTCG**TTTACA**GCTAGCTCAGTCCTAGG**TACAAT**ctcactatatcgccgatatagagcctactgagcacagctggctgagcacagctgtcaccggatgtgctttccggtctgatgagtccgtgaggacgaaacagcctctacaaataattttgtttaaactagaagtaagcgaggtacacatATG |
| P750 | CTCAGTGGCGCGCCTCAGTCCTCG**TTGACA**GCTAGCTCAGTCCTAGG**TACAAT**ctcactatatcgccgatatagagcctactgagcacagctggctgagcacagctgtcaccggatgtgctttccggtctgatgagtccgtgaggacgaaacagcctctacaaataattttgtttaaactagaagtaagcgaggtacacatATG |
| P1000 | AAAATATATTTTTCAAAAGTATCG**TTGACA**GCTAGCTCAGTCCTAGG**TACAAT**ctcactatatcgccgatatagagcctactgagcacagctggctgagcacagctgtcaccggatgtgctttccggtctgatgagtccgtgaggacgaaacagcctctacaaataattttgtttaaactagaagtaagcgaggtacacatATG |
| NAT | accgatccgcacaatgccagctcaagcctgagggtaaaccgcaagccaacatttgcgccgtttttaaacgaagatg**ttgacc**acactccttctcgccaa**tataCT**CAGTATATCGCAGATATAGAGCCTgcaattacagccatcgtacgagccctggctgagcacagctgtcaccggatgtgctttccggtctgatgagtccgtgaggacgaaacagcctctacaaataattttgtttaaactagaagtaagcgaggtacacatATG |
| WT | ggctgtcgacctttgaaaagttcg**tttaca**gctagctcagtcctagg**tacaat**CTCAGTATATCGCAGATATAGAGCCTactgagcacagctggctgagcacagctgtcaccggatgtgctttccggtctgatgagtccgtgaggacgaaacagcctctacaaataattttgtttaaactagaagtaagcgaggtacacatATG |
| INV | ggctgtcgacctttgaaaagttcg**tttaca**gctagctcagtcctagg**tacaat**AGGCTCTATATCTGCGATATACTGAGactgagcacagctggctgagcacagctgtcaccggatgtgctttccggtctgatgagtccgtgaggacgaaacagcctctacaaataattttgtttaaactagaagtaagcgaggtacacatATG |
| V2 | ggctgtcgacctttgaaaagttcg**tttaca**gctagctcagtcctagg**tacaat**CTCACTATATCGCAGATATAGAGCCTactgagcacagctggctgagcacagctgtcaccggatgtgctttccggtctgatgagtccgtgaggacgaaacagcctctacaaataattttgtttaaactagaagtaagcgaggtacacatATG |
| V3 | ggctgtcgacctttgaaaagttcg**tttaca**gctagctcagtcctagg**tacaat**CTCACTATATCGCCGATATAGAGCCTactgagcacagctggctgagcacagctgtcaccggatgtgctttccggtctgatgagtccgtgaggacgaaacagcctctacaaataattttgtttaaactagaagtaagcgaggtacacatATG |
| R1 | ggctgtcgacctttgaaaagttcg**tttaca**gctagctcagtcctagg**tacaat**ctcactatatcgccgatatagagcctactgagcacagctggctgagcacagctgtcaccggatgtgctttccggtctgatgagtccgtgaggacgaaacagcctctacaaataattttgtttaaactagGACTTTAAAGTCCAAacatATG |
| R2 | ggctgtcgacctttgaaaagttcg**tttaca**gctagctcagtcctagg**tacaat**ctcactatatcgccgatatagagcctactgagcacagctggctgagcacagctgtcaccggatgtgctttccggtctgatgagtccgtgaggacgaaacagcctctacaaataattttgtttaaactagTGAACCACGAGGCCTacatATG |
| R3 | ggctgtcgacctttgaaaagttcg**tttaca**gctagctcagtcctagg**tacaat**ctcactatatcgccgatatagagcctactgagcacagctggctgagcacagctgtcaccggatgtgctttccggtctgatgagtccgtgaggacgaaacagcctctacaaataattttgtttaaactagTTAGATAAGGAGGTTacatATG |
| R4 | ggctgtcgacctttgaaaagttcg**tttaca**gctagctcagtcctagg**tacaat**ctcactatatcgccgatatagagcctactgagcacagctggctgagcacagctgtcaccggatgtgctttccggtctgatgagtccgtgaggacgaaacagcctctacaaataattttgtttaaactagCGCCTAGAGGGGTTacatATG |
| R5 | ggctgtcgacctttgaaaagttcg**tttaca**gctagctcagtcctagg**tacaat**ctcactatatcgccgatatagagcctactgagcacagctggctgagcacagctgtcaccggatgtgctttccggtctgatgagtccgtgaggacgaaacagcctctacaaataattttgtttaaactagAAGTAAGCGAGGTACacatATG |

**Supplementary Table 2: Mutations in CamR variants evolved for alternative monoterpenes**

| Mutant number | Borneol | Fenchol | Eucalyptol | Camphene |
|---|---|---|---|---|
| 1 | Y58F, A128V | A139T | Y58F | Y58F |
| 2 | G67V, M111L | A139T | Y58F | Y58F |
| 3 | Y58F | A139T | Y58F | Y58F |
| 4 | Y43F, Y58F, Q125L | A139T | Y58F | Y58F, S110P, R137H |
| 5 | Y58F, D114V | A139T | Y58F | Y58F, A17V |
| 6 | Y58F, E70V | A139T | Y58F, T35S | Y58F, T69A |
| 7 | Y58F, G146S | Y58F | Y58F, A83T | A139T |
| 8 | Y58F | Y58F | A139T | A83T, A139S |
| 9 | Q47R, Y58F, T133S | Y58F, M104L | E70K, K76Q | H102N |
| 10 | Y58F, K75R, F126I | G67V, M111L | L75I | M111L, A120E, D121E, A139T |

**Supplementary Table 3: Mutations in two CamR generations evolved for fenchol**

| Variant number | Generation 2 (FEN2.X) | Generation 3 (FEN3.X) |
|---|---|---|
| 1 | (A139T) A83D | (A139T, G118R) V110I |
| 2 | (A139T) H102N **(x2)** | (A139T, G118R) M104L **(x3)** |
| 3 | (A139T) G118R | (A139T, F126Y) V110I |
| 4 | (A139T) F126Y | (A139T, H102N) R137H, Q164P |
| 5 | | (A139T, G118R) R157H |
| 6 | | (A139T, A83D) K76R, Y58F |

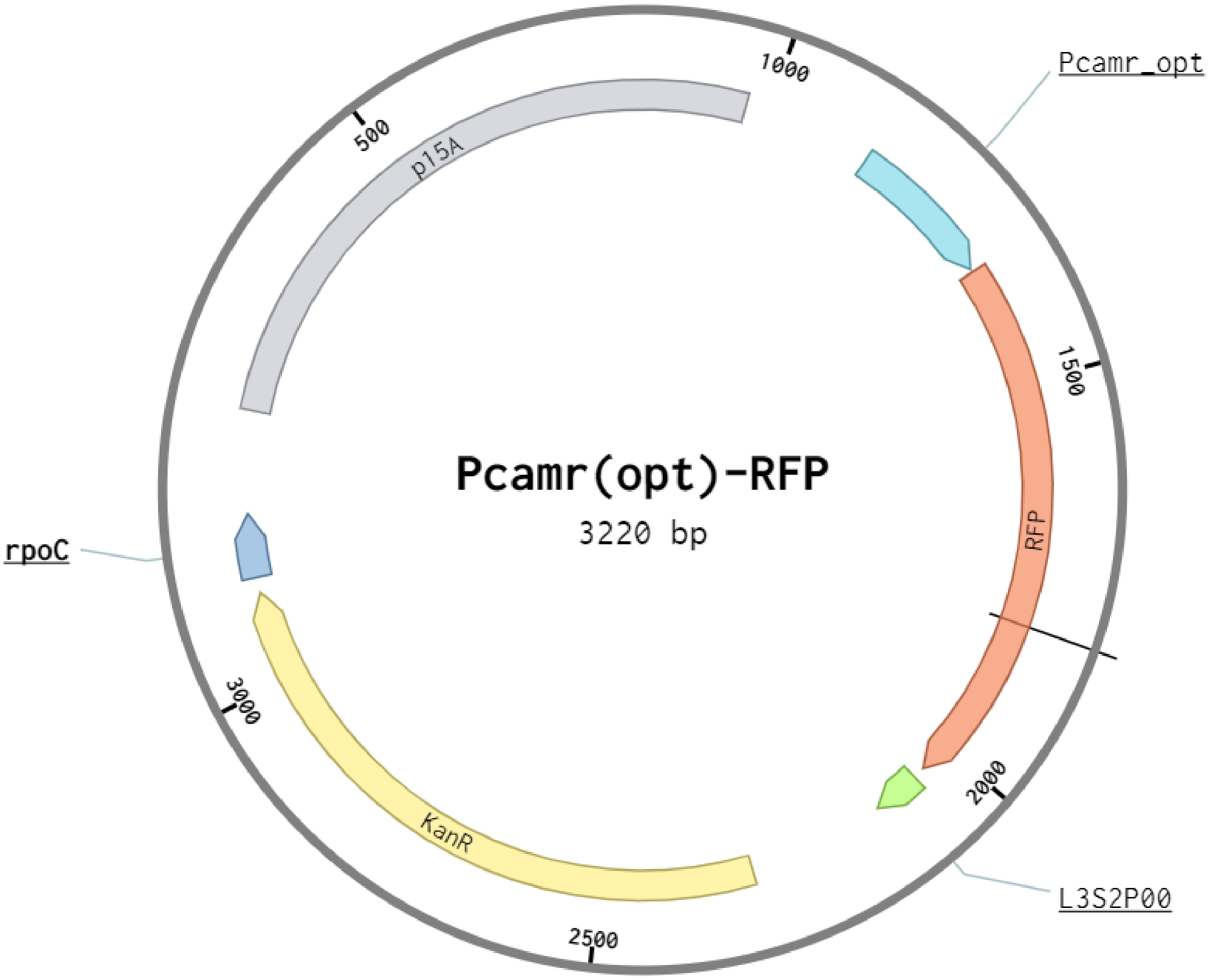


>Pcamr(opt)-RFP

cgttgaatatggctcccttaacgtgaggaagttcctatactttctagagaataggaacttctacagatggacttgggttggcggtttcaggagtaggtgctt
cctcgctcactgactcgctgcacgaggcagacctcagcgctagcggagtgtatactggcttactatgttggcactgatgagggtgtcagtgaagtgctt
catgtggcaggagaaaaaaggctgcaccggtgcgtcagcagaatatgtgatacaggatatattccgcttcctcgctcactgactcgctacgctcggtc
gttcgactgcggcgagcggaaatggcttacgaacggggcggagatttcctggaagatgccaggaagatacttaacagggaagtgagagggccgc
ggcaaagccgtttttccataggctccgcccccctgacaagcatcacgaaatctgacgctcaaatcagtggtggcgaaacccgacaggactataaaga
taccaggcgtttccccctggcggctccctcgtgcgctctcctgttcctgcctttcggtttaccggtgtcattccgctgttatggccgcgtttgtctcattcca
cgcctgacactcagttccgggtaggcagttcgctccaagctggactgtatgcacgaaccccccgttcagtccgaccgctgcgccttatccggtaacta
tcgtcttgagtccaacccggaaagacatgcaaaagcaccactggcagcagccactggtaattgatttagaggagttagtcttgaagtcatgcgccggtt
aaggctaaactgaaaggacaagttttggtgactgcgctcctccaagccagttacctcggttcaaagagttggtagctcagagaaccttcgaaaaaccg
ccctgcaaggcggttttttcgttttcagagcaagagattacgcgcagaccaaaacgatctcaagaagatcatcttattaaggggtctgacgctcagtgga
acgaaaccctgttggtcaggttttcgggaggtgtgcgtctcgcatccggaaaggtgtgataggtagcgcagcaaataaaacgaaaggctcagtcgaa
agactgggcctttcgttttatctgttgtttgtcggtgaacgctctcctcaacgggctgtcgacctttgaaaagttcgtttacagctagctcagtcctaggtac
aatctcactatatcgccgatatagagcctactgagcacagctggctgagcacagctgtcaccggatgtgctttccggtctgatgagtccgtgaggacga
aacagcctctacaaataattttgtttaaactagaagtaagcgaggtacacatatggtcagtaaaggagaagctgtaattaaagagtttatgcgcttcaaag
tgcatatggaaggttccatgaacggacatgagttcgagattgaaggtgaaggtgaaggacgcccctacgagggcactcaaactgcaaagttaaaggt
tacaaaaggaggacctctgccattttcgtgggacattctgagcccgcagtttatgtacggcagccgcgcgttcatcaaacatcccgctgacatcccaga
ctattacaaacaatctttccccgaaggctttaaatgggaacgcgtgatgaactttgaagatggtggcgctgtgactgtgacccaggacacttcattagaa
gatggaaccctgatttacaaggtaaagctgcgcggcaccaactttcccccctgacggacctgtaatgcagaaaaaaacaatgggttgggaggctagta
cagagcgtttataccctgaggacggtgtcttaaaaggagacatcaagatggcgttacgtcttaaggatggtggtcgctatttagctgacttcaagaccac
ttataaagcaaagaagcccgtccaaatgcctggagcttataacgttgaccgtaagttagacatcacctcacataacgaggattacacagttgtcgaaca
gtatgagcgctcagaaggccgtcattcgactggtggaatggacgaactgtataaataataatcctgcttcattcctcggtaccaaattccagaaaagag

gggagcgggaaaccgctcccctttttcgttttggtccctcgctgcagtgtcgtaaaaaccagttggggagacgctgtgaatcccgggcaagcaaaat
cgctggaagttcctatactttctagagaataggaacttctttctaaatacattcaaatatgtatccgctcatgagacaataaccctgataaatgcttcaataat
attgaaaaaggaagagtatgagccatattcaacgggaaacgtcttgctccaggccgcgattaaattccaacatggatgctgatttatatgggtataaatg
ggctcgcgataatgtcgggcaatcaggtgcgacaatctatcgattgtatgggaagcccgatgcgccagagttgtttctgaaacatggcaaaggtagcg
ttgccaatgatgttacagatgagatggtcagactaaactggctgacggaatttatgcctcttccgaccatcaagcattttatccgtactcctgatgatgcat
ggttactcaccactgcgatccccgggaaaacagcattccaggtattagaagaatatcctgattcaggtgaaaatattgttgatgcgctggcagtgttcct
gcgccggttgcattcgattcctgtttgtaattgtccttttaacagcgatcgcgtatttcgcctcgctcaggcgcaatcacgaatgaataacggtttggttgat
gcgagtgattttgatgacgagcgtaatggctggcctgttgaacaagtctggaaagaaatgcataagcttttgccattctcaccggattcagtcgtcactca
tggtgatttctcacttgataaccttatttttgacgaggggaaattaataggttgtattgatgttggacgagtcggaatcgcagaccgataccaggatcttgc
catcctatggaactgcctcggtgagttttctccttcattacagaaacggctttttcaaaaatatggtattgataatcctgatatgaataaattgcagtttcatttg
atgctcgatgagtttttctaagttgtgatggcggtaggaatgtaatcgttaatccgcaaataacgtaaaaacccgcttcggcgggtttttttatgggggga
gtttagggaaagagcatttgtcatcc

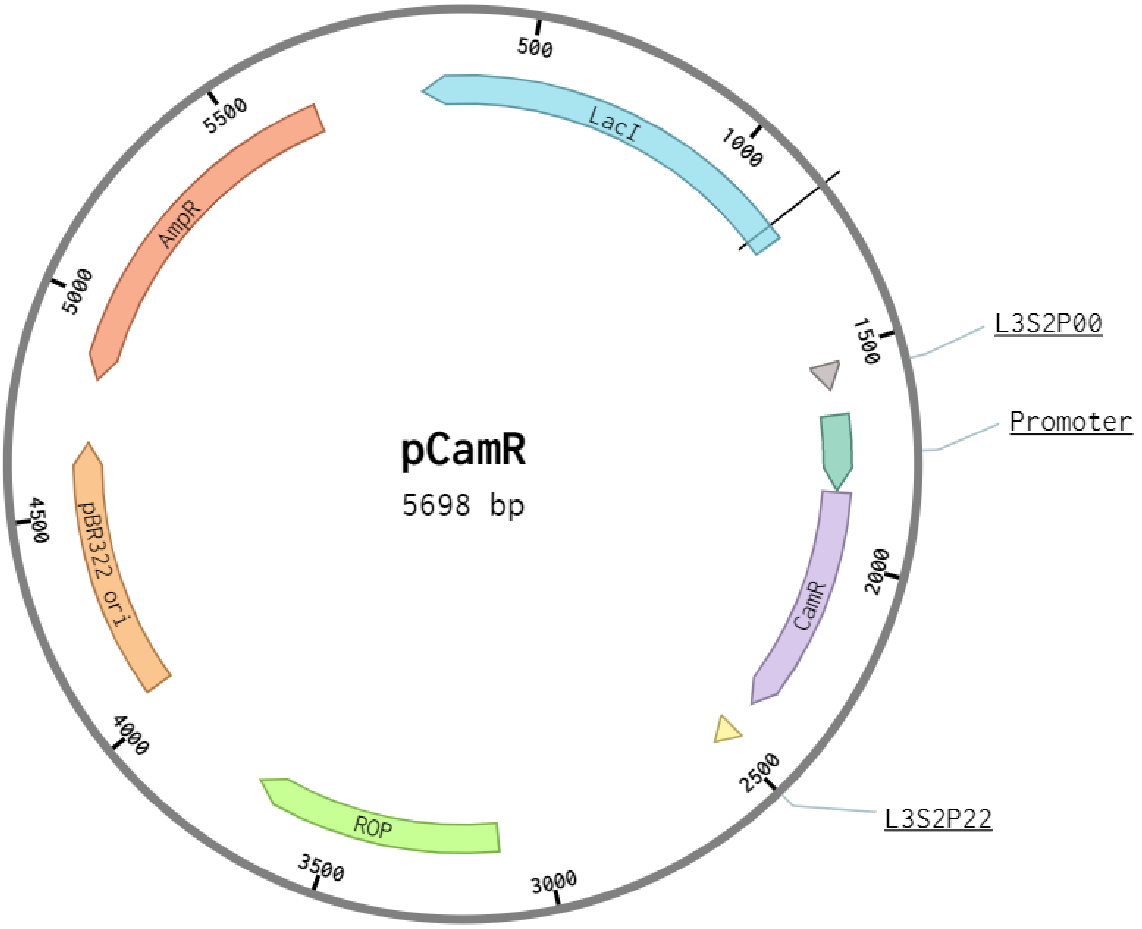


>pCamR

caatattattgaagcatttatcagggttattgtctcatgagcggatacatatttgaatgtatttagaaaaatatgcgccttgagcgacacgaattatgcagtg
atttacgacctgcacagccataccacagcttccgatggctgcctgacgccagaagcattggtgcaccgtgcagtcgatgataagctgtcaaacatgag
aattgtgcctaatgagtgagctaacttacattaattgcgttgcgctcactgcccgctttccagtcgggaaacctgtcgtgccagctgcattaatgaatcgg
ccaacgcgcggggagaggcggtttgcgtattgggcgccagggtggtttttcttttcaccagtgagacgggcaacagctgattgcccttcaccgcctgg
ccctgagagagttgcagcaagcggtccacgctggtttgccccagcaggcgaaaatcctgtttgatggtggttaacggcgggatataacatgagctatc
ttcggtatcgtcgtatcccactaccgagatatccgcaccaacgcgcagcccggactcggtaatggcgcgcattgcgcccagcgccatctgatcgttg
gcaaccagcatcgcagtgggaacgatgccctcattcagcatttgcatggtttgttgaaaaccggacatggcactccagtcgccttcccgttccgctatc
ggctgaatttgattgcgagtgagatatttatgccagccagccagacgcagacgcgccgagacagaacttaatgggcccgctaacagcgcgatttgct
ggtgacccaatgcgaccagatgctccacgcccagtcgcgtaccatcttcatgggagaaaataatactgttgatgggtgtctggtcagagacatcaaga
aataacgccggaacattagtgcaggcagcttccacagcaatggcatcctggtcatccagcggatagttaatgatcagcccactgacgcgttgcgcga
gaagattgtgcaccgccgctttacaggcttcgacgccgcttcgttctaccatcgacaccaccacgctggcacccagttgatcggcgcgagatttaatcg

ccgcgacaatttgcgacggcgcgtgcagggccagactggaggtggcaacgccaatcagcaacgactgtttgcccgccagttgttgtgccacgcggt
tgggaatgtaattcagctccgccatcgccgcttccactttttcccgcgttttcgcagaaacgtggctggcctggttcaccacgcgggaaacggtctgata
agagacaccggcatactctgcgacatcgtataacgttactggtttcacattcaccaccctgaattgactctcttccgggcgctatcatgccataccgcga
aaggttttgcaccattcgatggtgtcgggacgtcaggtggcacttttcggggaaatgtgcgcggaacccctatttgttgcggccgcgaagacagcgtt
atcagagatgagacaccgtgcgataatgtcgggcaatcaggtgcgactcggtaccaaattccagaaaagaggggagcgggaaaccgctcccctttt
tcgttttggtcccaatctatcgattgtatggacttcatcttcattacctcgtatcattgtacacctgccgaacgcaagggcatgggctgtcgacctttgaaaa
gtacctttacggctagctcagtcctaggtatagtgctagcaatcaagatactgagcacagctgtcaccggatgtgctttccggtctgatgagtccgtgag
gacgaaacagcctctacaaataattttgtttaaactagctgccgcagacccgcacatatggacattaagcagtcgttacttcacgcagctatgcgcttact
tagcgccaagggccgcgacggagctaccatgcgtgcaatctgcgccgaagtgggtgttactcctccaactctgtatcaccattatggcgatcttcaag
gtttacataaagcagctatcgatgaaacataccgtcaggtcgcagaggcatatcacggcggaaccgaggagcgtggaccgttaaaaggcatccgcg
acggctgggccacattcctgcagttcgcctattcagagccaaatatgtgccgtatgttggttcaacatatcatggcaggagaaccccttccatggtgg
ctgataccccttcgcggagtcgcagatgatttagctcagttccatgcacaaggccgcttaaccttcccaccgcgtgaggccgcccaattattatggatgg
gcgctttgggtgcactgacctatgcactttcacgtgaaggagcaggttacactcaggacttggcactgcagaaggctaaattagatattacacttgtagc
actgtttaatatcgaggaggaataataatcctaagaattcgaggagtgcaggctcggtaacatacggtctagctatctgactatcgccgctgtgagctcg
gtaccaaattccagaaaagaggccgcgaaagcggcctttttcgttttggtccaagcccgatgcgccagagttgtttctgaaacatggcaaaggtatca
ctagtcttcgcggccgccatgctgtccaggcaggtagatgacgaccatcagggacagcttcaaggatcgctcgcggctcttaccagcctaacttcgat
cattggaccgctgatcgtcacggcgatttatgccgcctcggcgagcacatggaacgggttggcatggattgtaggcgccgccctataccttgtctgcc
tccccgcgttgcgtcgcggtgcatggagccgggccacctcgacctgaatggaagccggcggcacctcgctaacggattcaccactccaagaattgg
agccaatcaattcttgcggagaactgtgaatgcgcaaaccaacccttggcagaacatatccatcgcgtccgccatctccagcagccgcacgcggcg
catctcgggcagcgttgggtcctggccacgggtgcgcatgatcgtgctcctgtcgttgaggacccggctaggctggcggggttgccttactggttagc
agaatgaatcaccgatacgcgagcgaacgtgaagcgactgctgctgcaaaacgtctgcgacctgagcaacaacatgaatggtcatcggtttccgtgtt
tcgtaaagtctggaaacgcggaagtcagcgccctgcaccattatgttccggatctgcatcgcaggatgctgctggctaccctgtggaacacctacatct
gtattaacgaagcgctggcattgaccctgagtgatttttctctggtcccgccgcatccataccgccagttgtttaccctcacaacgttccagtaaccgggc
atgttcatcatcagtaacccgtatcgtgagcatcctctctcgtttcatcggtatcattacccccatgaacagaaatcccccttacacggaggcatcagtga
ccaaacaggaaaaaaccgcccttaacatggcccgctttatcagaagccagacattaacgcttctggagaaactcaacgagctggacgcggatgaac
aggcagacatctgtgaatcgcttcacgaccacgctgatgagctttaccgcagctgcctcgcgcgtttcggtgatgacggtgaaaacctctgacacatg
cagctcccggagacggtcacagcttgtctgtaagcggatgccgggagcagacaagcccgtcagggcgcgtcagcgggtgttggcgggtgtcggg
gcgcagccatgacccagtcacgtagcgatagcggagtgtatactggcttaactatgcggcatcagagcagattgtactgagagtgcaccggtgtgaa
ataccgcacagatgcgtaaggagaaaataccgcatcaggcgctcttccgcttcctcgctcactgactcgctgcgctcggtcgttcggctgcggcgag
cggtatcagctcactcaaaggcggtaatacggttatccacagaatcaggggataacgcaggaaagaacatgtgagcaaaaggccagcaaaaggcc
aggaaccgtaaaaaggccgcgttgctggcgtttttccataggctccgcccccctgacgagcatcacaaaaatcgacgctcaagtcagaggtggcga
aacccgacaggactataaagataccaggcgtttccccctggaagctccctcgtgcgctctcctgttccgaccctgccgcttaccggatacctgtccgc
ctttctcccttcgggaagcgtggcgctttctcatagctcacgctgtaggtatctcagttcggtgtaggtcgttcgctccaagctgggctgtgtgcacgaac
cccccgttcagcccgaccgctgcgccttatccggtaactatcgtcttgagtccaacccggtaagacacgacttatcgccactggcagcagccactggt
aacaggattagcagagcgaggtatgtaggcggtgctacagagttcttgaagtggtggcctaactacggctacactagaaggacagtatttggtatctg
cgctctgctgaagccagttaccttcggaaaaagagttggtagctcttgatccggcaaacaaaccaccgctggtagcggtggtttttttgtttgcaagcag
cagattacgcgcagaaaaaaaggatctcaagaagatcctttgatcttttctacggggtctgacgctcagtggaacgaaaactcacgttaagggattttgg
tcatgagattatcaaaaaggatcttcacctagatccttttaaattaaaaatgaagttttaaatcaatctaaagtatatatgagtaaacttggtctgacagttacc
aatgcttaatcagtgaggcacctatctcagcgatctgtctatttcgttcatccatagttgcctgactccccgtcgtgtagataactacgatacgggagggct
taccatctggccccagtgctgcaatgataccgcgcgatccacgctcaccggctccagatttatcagcaataaaccagccagccggaagggccgagc
gcagaagtggtcctgcaactttatccgcctccatccagtctattaattgttgccgggaagctagagtaagtagttcgccagttaatagtttgcgcaacgtt
gttgccattgctgcaggcatcgtggtgtcacgctcgtcgtttggtatggcttcattcagctccggttcccaacgatcaaggcgagttacatgatcccccat
gttgtgcaaaaaagcggttagctccttcggtcctccgatcgttgtcagaagtaagttggccgcagtgttatcactcatggttatggcagcactgcataatt
ctcttactgtcatgccatccgtaagatgcttttctgtgactggtgagtactcaaccaagtcattctgagaatagtgtatgcggcgaccgagttgctcttgcc
cggcgtcaacacgggataataccgcgccacatagcagaactttaaaagtgctcatcattggaaaacgttcttcggggcgaaaactctcaaggatctta
ccgctgttgagatccagttcgatgtaacccactcgtgcacccaactgatcttcagcatcttttactttcaccagcgtttctgggtgagcaaaaacaggaag
gcaaaatgccgcaaaaaagggaataagggcgacacggaaatgttgaatactcatactcttcctttt

**Supplementary Table 4.** Relevant plasmid maps and sequences
